# MinorityReport, software for generalized analysis of causal genetic variants

**DOI:** 10.1101/098731

**Authors:** Jeremy A Horst, Wesley Wu, Joseph L DeRisi

**Affiliations:** Department of Biochemistry and Biophysics, University of California San Francisco School of Medicine; Howard Hughes Medical Institute, 1700 4th St. QB3 Room 404 San Francisco CA 94158-2330

**Keywords:** Bioinformatics, mutation, genotype-phenoype, drug mechanism, genetic etiology, CNV, missense

## Abstract

**Background:** The widespread availability of next generation genome sequencing technologies has enabled a wide range of variant detection applications, especially in cancer and inborn genetic disorders. For model systems and microorganisms, the same technology may be used to discover the causative mutations for any phenotype, including those generated in response to chemical perturbation. In the case of pathogenic organisms, these approaches have allowed the determination of drug targets by means of resistance selection followed by genome sequencing.

**Results:** Here, we present open source software written in python, MinorityReport, to facilitate the comparison of any two sets of genome alignments for the purpose of rapidly identifying the spectrum of nonsynonymous changes, insertions or deletions, and copy number variations in a presumed mutant relative to its parent. Specifically, MinorityReport relates mapped sequence reads in SAM format output from any alignment tool for both the mutant and parent genome, relative to a reference genome, and produces the set of variants that distinguishes the mutant from the parent, all presented in an intuitive, straightforward report format. MinorityReport features tunable parameters for evaluating evidence and a scoring system that prioritizes reported variants based on relative proportions of read counts supporting the variant in the mutant versus parent data sets. We demonstrate the utility of MinorityReport using publicly available data sets that we previously published to find the determinants of resistance for novel anti-malarial drugs.

**Conclusions:** MinorityReport is readily available (github: xxxxxxx) to identify the genetic mechanisms of drug resistance in plasmodium, genotype-phenotype relationships in human diads, or genomic variations between any two related organisms.

## BACKGROUND

### Sequencing to characterize genotype-phenotype relationships

The advent of robust high throughput genetic sequencing methods has made genome sequencing tractable for any sample from any organism. Comparison of genomes for closely related organisms with different physiologic, behavioral, or anatomic phenotypes has been increasingly used to characterize the genetic determinants of the feature, i.e. genotype-phenotype relationships. Differences in the genome sequences of 2 related organisms with a different phenotype describe the possible causal genetic variants (see Figure 1). We have used this technique to find the molecular determinants of resistance for novel compounds against the causative agent of the most deadly form of malaria, *P. falciparum* [1], including a drug now in clinical trials [2], and an inhibitor of the apicoplast – an organelle unique to this class of organisms [3]. We used the same principles to find novel mutant genes underlying human craniofacial congenital malformations including craniosynostosis [4], auriculocondylar syndrome and thereby the universal signal for lower jaw patterning [5], and acromelic frontonasal dysostosis [6]. While we have focused on congenital anomalies and drug targets, this experimental system is relevant to any genotype-phenotype study.

**Figure 1.**
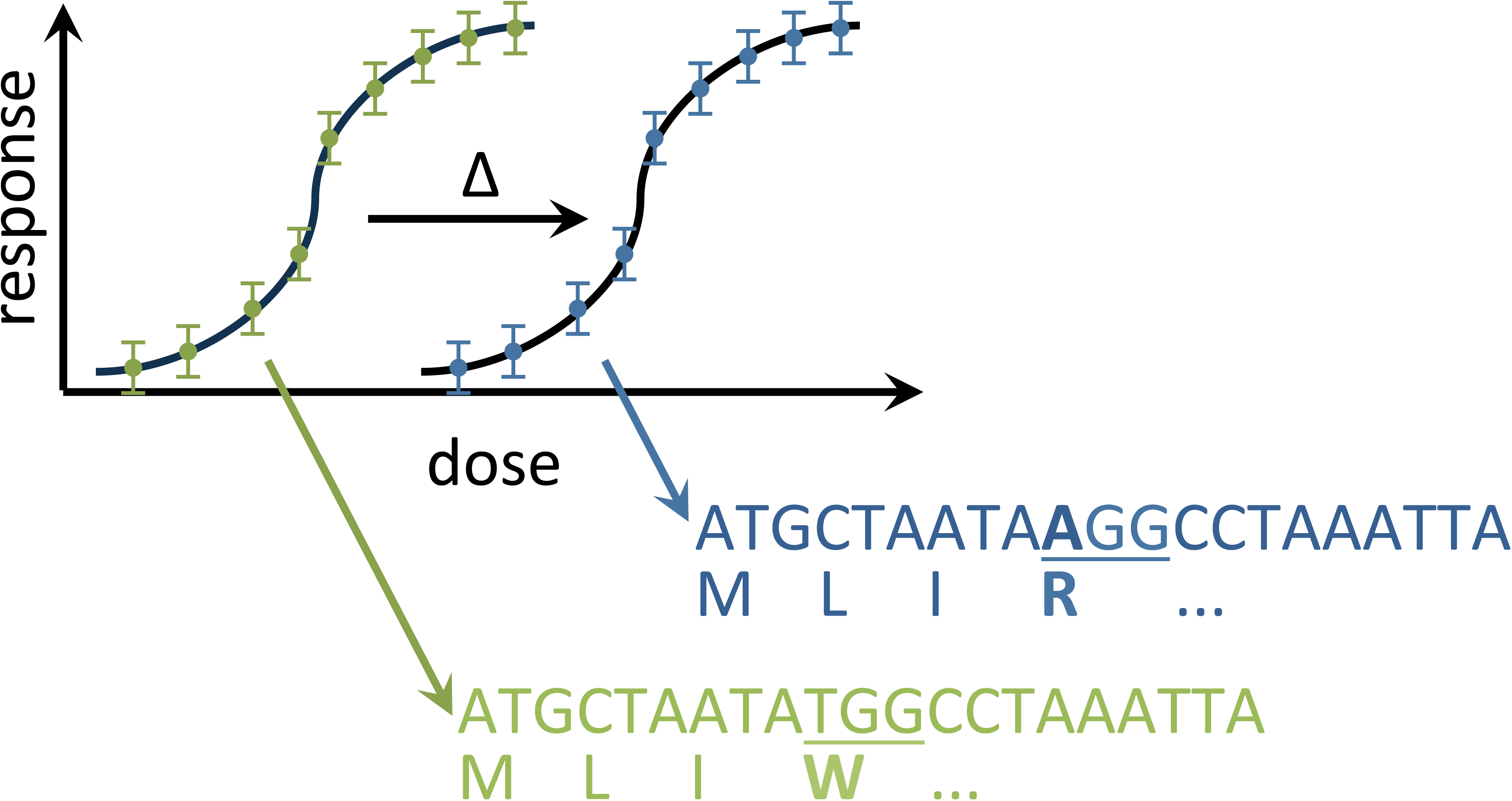
Experimental scheme. MinorityReport deciphers nonsynonymous mutations, including those that enable drug resistance in pathogens. A susceptible pathogen strain (green) is cultivated in sub-LD100 concentrations of a drug until a resistant strain (blue) emerges. The genomes are sequenced, aligned to a reference genome, and MinorityReport outputs the causative mutation, which is often also the drug target.

### How to

To perform this analysis, genomic sequencing reads from each organism (e.g. in FASTQ format) are first aligned to the reference genome FASTA file, resulting in Sequence Alignment / Map (SAM) format files [7]. The observed variations from the reference genome are then evaluated for sufficient supporting evidence by quantity and proportion of reads in the mutant, and support for non-variation in the parent. The variants that pass these threshold filters are then placed in the context of coding sequences, defined in gene file format files (GFF3), to determine whether the variant changes the gene product, *i.e.* is nonsynonymous (Figure 2). Variants observed consistently in multiple independent pairs of organisms that differ in the same manner are likely to be the causal genetic variant that underlies the phenotype. In the case of drug resistance selections, the underlying assumption for a mutation or copy-number variant which leads to resistance is that it will be under purifying selection unlike other variants present in the population at the time of the initial selection.

**Figure 2.**
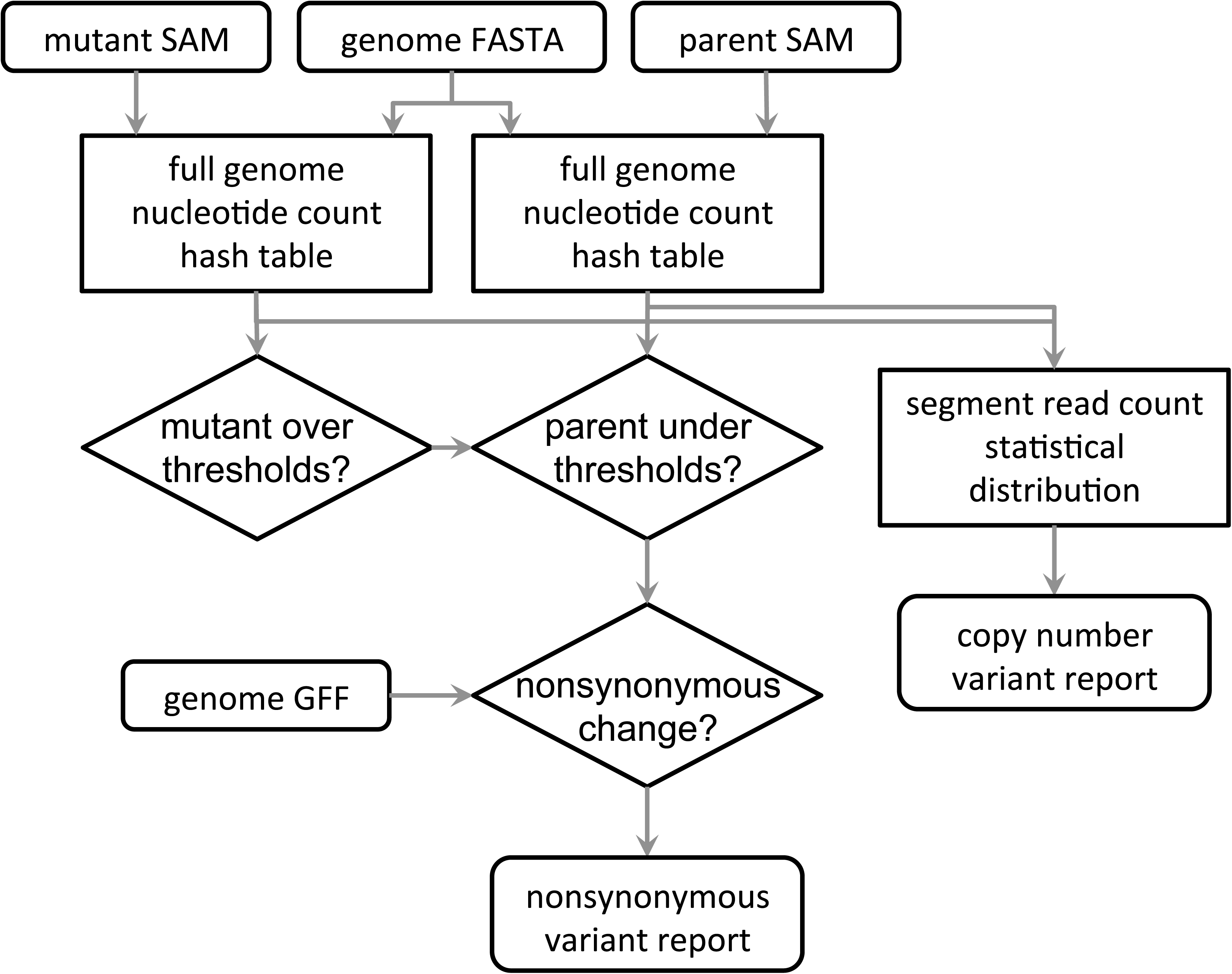
Software flow chart. MinorityReport relates mapped sequence reads in SAM format output from any alignment tool for both the mutant and parent genome, relative to a reference genome, to the protein coding sequence, and produces the set of nonsynonymous variant that distinguish the mutant from the parent.

This process requires the reference genome and gene model, and deep sequencing data for unaffected parents and the affected child for humans, or similarly the drug-sensitive parent strain and the derivative drug-resistant mutants for pathogens. The reference genome is required as a contextual template to align the sequencing reads and thereby identify genetic variants. The gene model is needed to identify whether a variant is in a protein-coding region, and whether the variant changes the resulting protein sequence.

While this process is straightforward in concept, many bioinformatic analysis computer programs work only on one or an arbitrary set of organisms, e.g. human and mouse, or do not integrate both mutation and copy-number variant analysis into a unified pipeline. A few robust generalized tools are available to call variants between one set of sequencing reads against any reference genome, and a subset of these call nonsynonymous variants. However, these tools require extensive data preparation, have software library dependencies that complicate installation, are sensitive to the many imperfections in gene models for the many poorly characterized organisms (i.e. they fail to complete running or give any output), and they do not perform comparative analysis between multiple sequencing sets. In practice, many researchers cobble together pipelines based on several different tools to accomplish this task.

### Freedom of alignment

Alignment algorithms continue to evolve at rapid pace, and the most recent iterations capture alignments to much more dissimilar sequences with inexact matches and significant insertions or deletions (indels). e.g. BowTie2, handle indels of limited size, and further sequence differences [8]. Newer algorithms such as STAR [9] and GSNAP [10] handle extensive indels, and highly distinct but still recognizably similar read-genome sequence matches. These advances are useful to address the genotype-phenotype problem. Fortunately, these and other genome alignment algorithms have converged on a standard SAM format [7] which enable downstream applications, such as the one described here, to be independent of the base aligner.

### CNVs

Another major type of genetic anomaly that can underlie genotype-phenotype relationships is a gene copy number variant (CNV). When the function of a gene becomes particularly adaptive, useless, or harmful, environmental pressures can select for the replication or loss of any number of copies of a gene or set of genes [11]. Because CNVs affect gene dosage, they have the potential for physiologic impact. Various approaches are used to detect CNVs on a large scale, with the primary clinical tool being array comparative genomic hybridization [12]. Algorithms have been constructed to use indirect evidence to evaluate CNVs, such as inference through analysis of single nucleotide polymorphism (SNP) chips [13]. However, as with all microarray technologies, next generation genome sequencing technologies are replacing the use of array CGH and SNP chips due to their portability to any biological system, and unbiased sequence data generation. The normalized abundance of sequencing reads that map to each gene is a direct measure of the number of copies of the gene. CNVs can be estimated using sparse read coverage data sets as well [11]. However, the enormous number of reads accessible through contemporary sequencing instruments (e.g. the Illumina HiSeq series) makes direct assessment through deep coverage straightforward.

## RESULTS AND DISCUSSION

Here, we present software, MinorityReport, to facilitate the comparison of any two sets of genome alignments for the purpose of rapidly identifying the spectrum of nonsynonymous or indel changes in a presumed mutant relative to its parent. In addition, the data model we use for detecting nonsynonymous variants integrates rapid assessment of CNVs. Tunable parameters for both analyses enable reporting of high or low purity nonsynonymous mutations, and large or small range CNV detection. This software does not address noncoding variants such as promoter or enhancer element mutations, intergenic variants that may occur in encoded functional RNAs, nor chromosomal rearrangements, which may be assessed with the same sequencing data. In this manuscript we elaborate the algorithms deployed in this software, and demonstrate use on freely available data sets that carry the genetic determinants of resistance for anti-malarial drugs.

### Nonsynonymous variant caller algorithm

A simplified flow chart of the nonsynonymous variant caller algorithm in MinorityReport is shown in Figure 2. FASTA and GFF3 files for the reference genome are taken as input to build the gene model. Indices for genes, exons, and splice variants are taken from coding sequence lines (those with CDS entered in the phase column). Gene descriptions are taken from gene lines. Reverse compliment indices are translated for genes present on the negative strand (python list indices are inclusive at the beginning but exclusive at the end, so must be shifted 1 nucleotide towards the 3’ direction). Sequences are found from these indices in the reference genome FASTA file, and checked for start and stop codons. Next, a hash table (python dictionary) is made from the reference genome FASTA file to store the sequence evidence: for each sequence position in each chromosome a key is created for each of the possible 4 nucleotides, and a zero value is entered (indels are added later).

SAM files of sequencing reads aligned to the reference genome for the parent and mutant are each taken as input and applied into separate sequence evidence hash tables. For each mapped read pair, the mate pair read is checked for matched mapping location. The CIGAR entry is read for presence of insertions or deletions (indels) in the read. The mapped strand is extracted from the binary flag entry. Finally, each nucleotide in each paired read is added into the sequence evidence hash table. Insertions and deletions are added to the position entry as they occur, and counted the same way as nucleotides. An example position: “sequence_evidence[‘chr7’ ][ 1423712 ] = {‘A’:3, ‘C’:5, ‘G’:51, ‘T’:3, ‘–’:426, ‘AACTAC’:20}” where chr7 is the chromosome, 1423712 is the position on chromosome 7, and 3 A’s, 5 C’s, 51 G’s, 3 T’s, 426 deletions, and 20 ACTAC insertions are observed at the position. Insertions are denoted in the direction of the positive strand; the first nucleotide in the insertion entry is technically not part of the insertion, but is counted in the insertion to enable a consistent notation system.

Next, the sequence evidence hash tables are scanned for nonsynonymous variants that appear in the mutant but not the parent. Chromosomes are read in turn. Then evidence thresholds are applied to each entry of the position in a progression to maximize efficiency: variant of the reference genome, supra-threshold number of mutant reads and fraction of reads at the position support the variant, supra-threshold number of parent read coverage, subthreshold number of parent reads and fraction of reads at the position support the variant, the variant is in a coding sequence region, and the variant changes an amino acid in the translated protein sequence (nonsynonymous). Variants passing these criteria are reported (Table). Each reported variant is assigned a priority score as follows:

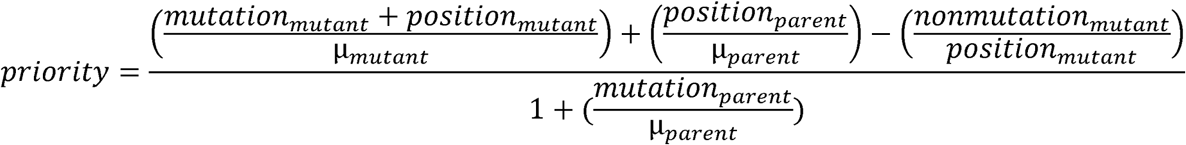

**Table.**
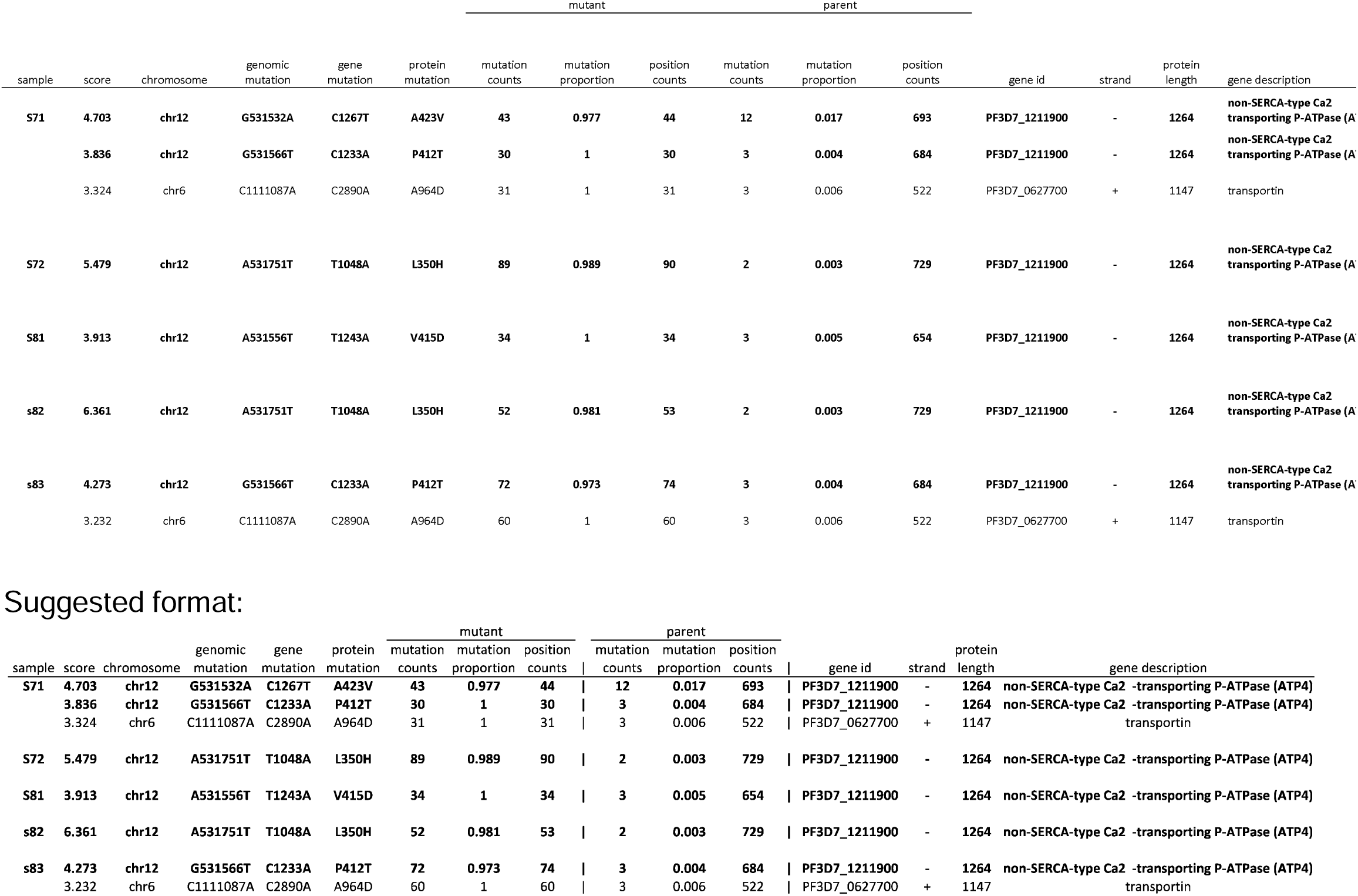
Sample MinorityReport output. MinorityReport output can be combined across multiple strains (s71, s72, s81, s82, s83) derived from the same parent strain in independent resistance experiments. The priority score aids identification of variants that reach near purity in mutant strains relative to the parent strain, with strong supporting evidence. The gene containing a mutation in all strains, here ATPase4, is the known determinant of resistance.

*mutation_mutant_* refers to the quantity of sequencing reads that contain the variant in the mutant sample, and similarly *mutation_parent_* refers to the same in the parent sample. *nonmutation _mutant_* refers to the quantity of reads that contain anything other than the variant in the mutant sample. *position* variables refer to the total number of reads overlapping the position of the variant in each sample, and μ refers to the mean number of reads overlapping any position in the genome for each sample.

### CNV caller algorithm

As each position in each chromosome is scanned for nonsynonymous variants unique to the mutant, the total counts in each of parent and mutant for each position are applied to a CNV evidence model. After the nonsynonymous variant scan is complete for each chromosome, the genome is evaluated for regions with significant differences in median read coverage between mutant and parent. The probability of the parent or mutant being significantly high or low in copy number (gene abundance) is estimated by relating the number of reads that map to the position to the mean and standard deviation for the chromosome, i.e. the z-score. Positions are considered significant when a number of reads are mapped in the mutant that correspond to being above or below a probability threshold or number of copies, with respect to the parent.

We calculate CNVs as the ratio of reads that map to each sliding window in the mutant and parent data sets, normalized by the total number of reads for the parent and mutant. We estimate the probability of observing a CNV at random for each variant by placing the log2 transform in the context of log2 normalized read ratios for all sliding windows assessed across the genome. The resulting z-score (z = (x − mean) / standard deviation) is transformed to a probability estimate by the Gaussian transformation.

The CNV-seq algorithm is widely used to calculate CNVs from sequencing data [11]. The CNV-seq algorithm calculates the probability for each observed CNV occurring at random using the covariation of the read numbers in the parent and mutant data sets, with the Geary-Hinkley transformation. As seen in Figures 3–5, read coverage is not homogenous. Neither does it follow a normal distribution. Areas with deviations in coverage due to external processing factors, such as the non-uniform distribution of so-called “random” hexamer-nucleotide primers, or the inaccuracy of genome alignments, may become inflated or diminished using this approach. The total amounts of reads in each set are also the result of processing factors. Thus, we examine the statistical probability of observing read ratios normalized by the total read count using a Gaussian distribution to the entire read set, i.e. including all chromosomes.

**Figure 3.**
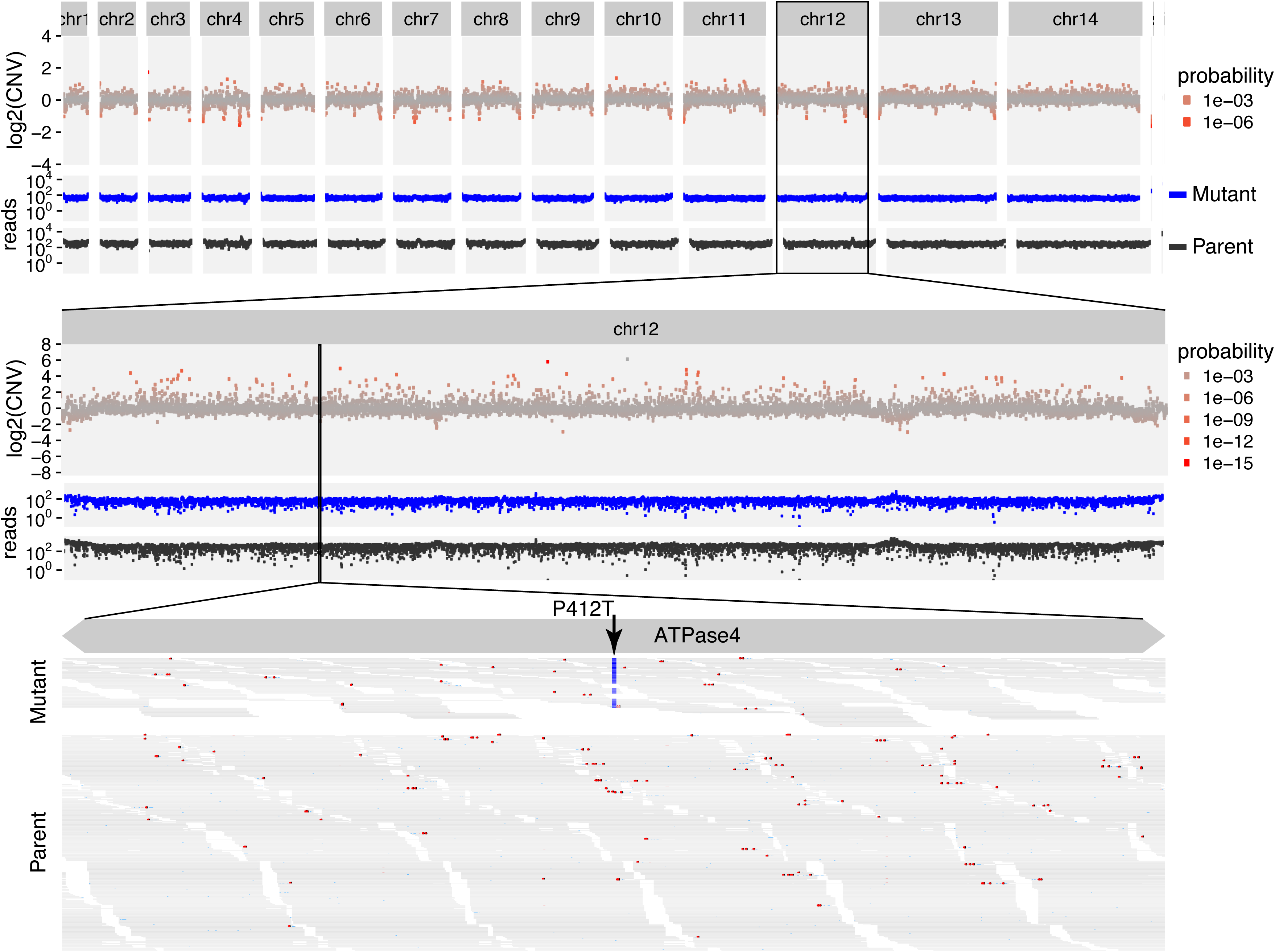
Nonsynonymous and copy number variant analyses to determine the drug target of antimalarial drug SJ733. The relative copy number for 3000 nucleotide tiles across the entire *P. falciparum* genome are mapped across all chromosomes (top panel). The read support for susceptible parent (black) and resistant mutant (blue) strains show a relatively even distribution of sequencing reads. Zooming into a reanalysis of chromosome 12 with 300 nucleotide tiles (middle panel) shows spurious copy number data in areas of relatively low read coverage. Zooming further in (bottom panel) shows that the variation that encodes P412T in ATPase4 (arrow) is present in 72 of 74 reads in the resistant mutant strain, but only 3 in 684 of the susceptible parent strain. This nonsynonymous mutation determines resistance to SJ733. Each panel is to scale.

### Application to determine target of novel anti-malarial compounds

#### SJ733

We recently reported the discovery of a novel antimalarial agent [2], which is now being tested for safety in humans. A key milestone for understanding the mechanism of action for SJ733 and its derivatives was to identify molecular determinants of resistance. We and others selected resistant sub-clones of a susceptible parent strain by long incubations in concentrations of the compound that allowed very few organisms to live, and identified the resistance loci by comparing genome sequence data between parent and resistant sub-clones (mutants). Comparing genomic sequencing reads between parent and mutants revealed a few variants in most mutants, while only one gene contained variants in all mutants: the gene encoding PfATP4, a sodium-dependent ATPase transporter [2].

We illustrate the process of simultaneously assessing copy number and nonsynonymous variation in the search for the mutation that enables resistance to SJ733 in Figure 3. For a haploid genome under drug selection, the underlying genetic assumption is that the causal mutant allele will be driven to purity in the population of surviving cells, and thus it is reasonable to set an absolute threshold of at least 90% for the proportion of reads that are variant at any given position. This threshold is a tunable parameter and would clearly need to be below 50% for a diploid genome, for example.

The sequencing data for this experiment are available through the NCBI Short Read Archive (SRA): SRX644366 is the mutant S83, SRX643297 is the parent strain, and other resistant strains derived in parallel independent experiments can be found through SRP043648.

The list of nonsynonymous variants found by MinorityReport (Table) demonstrates the coincident emergence of mutations in the same gene from 5 independent selection experiments. In fact, in 3 of 5 selections, the only mutation that passes these basic thresholds are those in PfATP4 [2]. Note that relaxing the mutant proportion threshold would reveal several high scoring nonsynonymous variants, but essentially all of these belong to highly repeated gene families, such as the PfEMP1 family of antigenic variation genes, which are inherently prone to spurious alignment. The appearance of these non-causal variants, such as transportin, in the data set exemplifies the rare occurrence of real or artifact passenger mutations that may be reported, and also emphasizes the utility of analyzing several independent selections.

Also, the CNV assessments differ between the granular genome-wide scan performed with 3000 nucleotide-long tiles (top) versus the chromosome-specific scan performed with 300 nucleotide-long tiles (middle); while this was done to construct an illustrative figure with a reasonable amount of data, it shows that tile size affects the appearance of copy number and likelihood calculations: larger tiles are recommended for more relevant probability values. In the case of PfATP4, our analysis produced no evidence of copy number variation associated with resistance to the compound.

#### MMV-08138

We recently reported the identification of the mechanism of action of a *Plasmodium* apicoplast inhibitor mMv-08138 [3]. As with SJ733, we compared the genome sequence of sub-clones raised for resistance to the drug against the susceptible parent strain. We demonstrate the emergence of a single nonsynonymous variation (IspD L244I) not observed in the parent strain, which reaches nearly complete (98.9%) purity in the resistant mutant shown in Figure 4. With the default threshold settings, 23 putative variants are output for this sample by MinorityReport (Supplementary Table 1). As well, the causal mutant receives over twice the priority score of any other variant.

**Figure 4.**
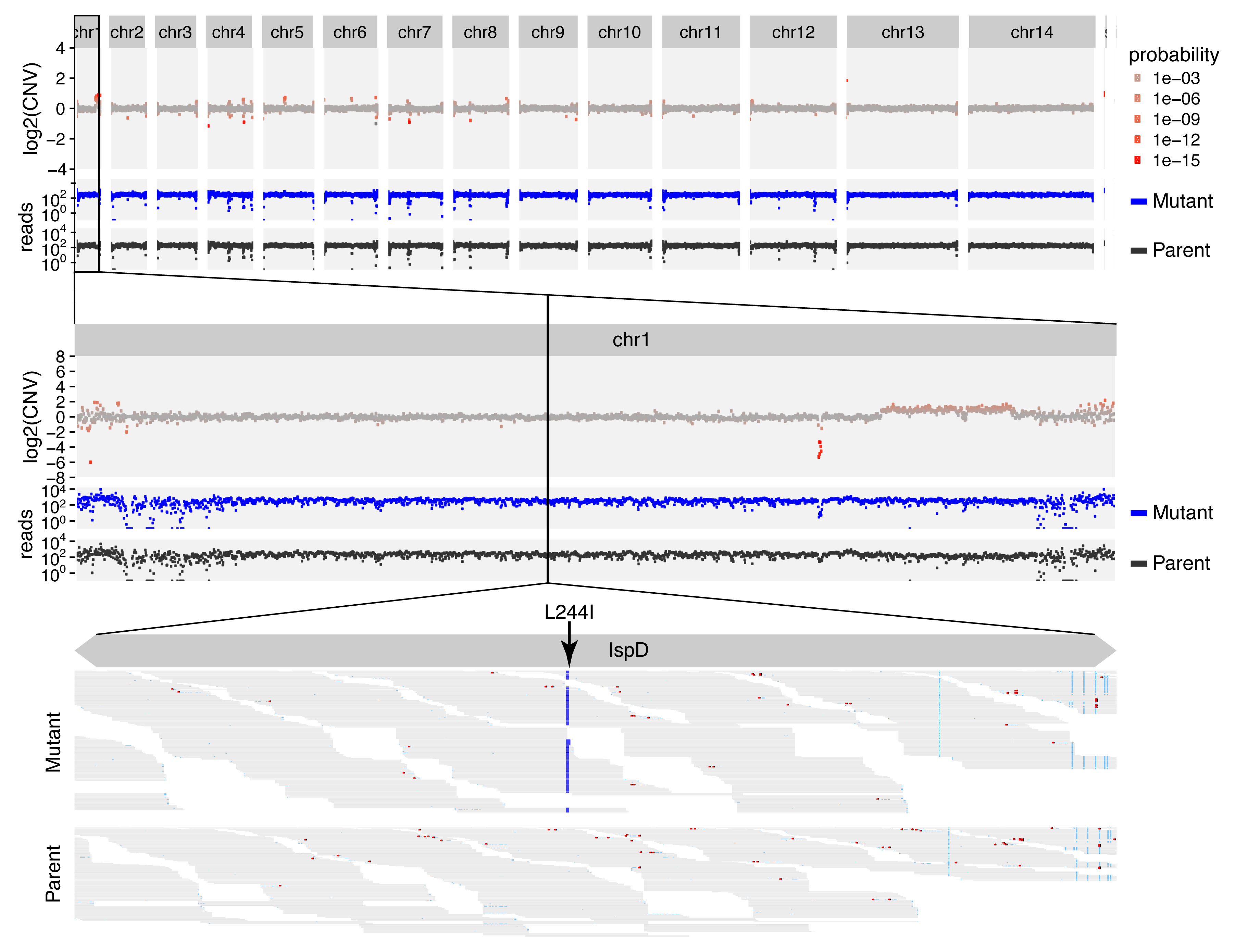
Nonsynonymous and copy number variant analyses to determine the drug target of apicoplast-specific inhibitor MMV-08138. Depictions match Figure 3. These data show no significant copy number variation. The bottom panel shows that the variation that encodes L244I in IspD (arrow) is present in 358 of 362 overlapping reads from the resistant mutant strain, but absent in all 242 from the susceptible parent strain. This nonsynonymous mutation determines resistance to MMV-08138.

#### DSM1

We also previously demonstrated that *P. falciparum* can achieve drug resistance to the dihydroorotate dehydrogenase inhibitor, DSM1, and perhaps generally, by randomly amplifying a large AT-repeat flanked region of the genome, then specifically further amplifying the region in progeny that successfully evade the drug effects [1]. Amplified regions likely contain the drug target, which can also contain nonsynonymous mutations that further aid resistance. MinorityReport successfully identified the amplified region (Figure 5), and 4 nonsynonymous mutations (Supplementary Table 2) that were previously reported.

**Figure 5.**
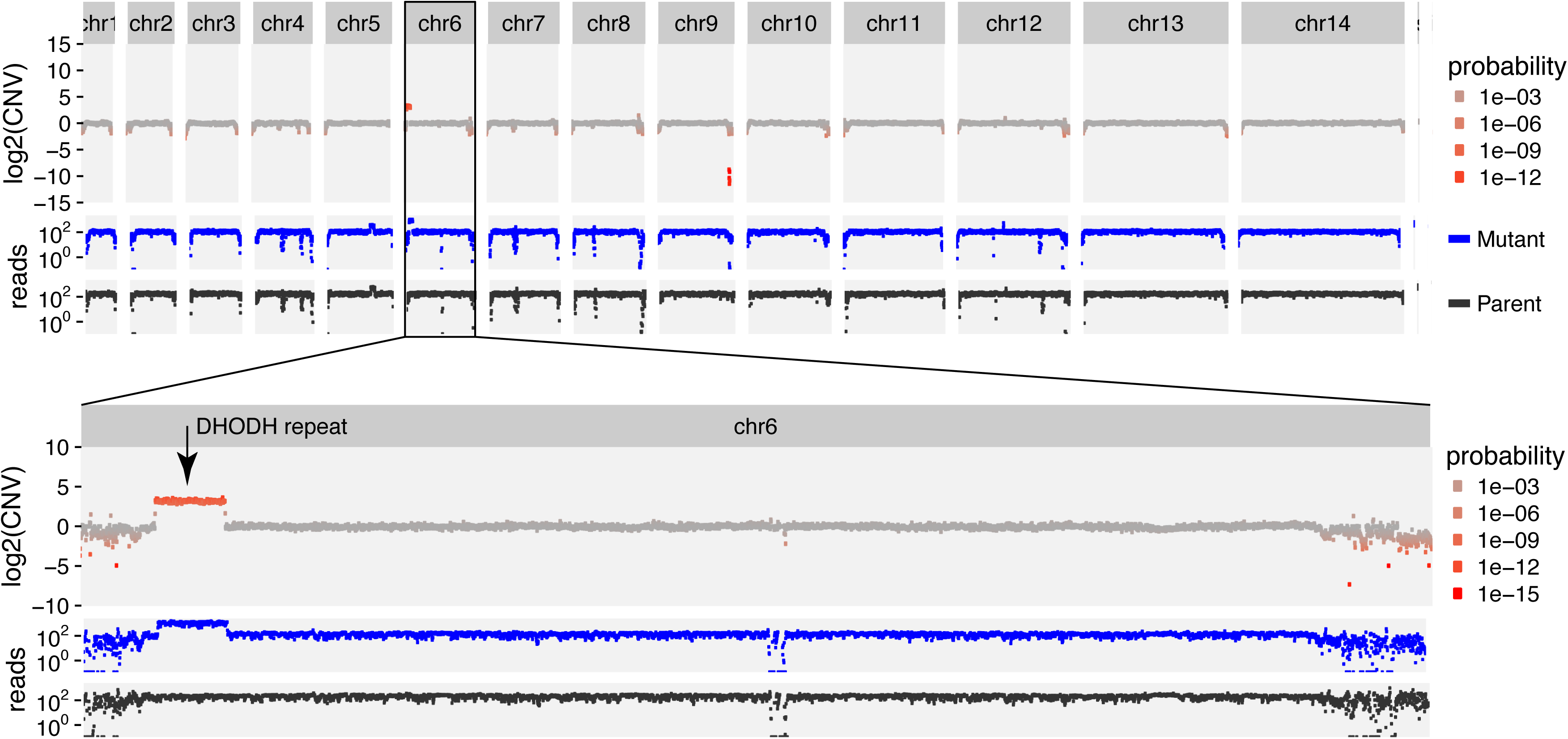
Nonsynonymous and copy number variant analyses to determine the drug target of antimalarial compound DSM1. Depictions match Figure 3. These data show a significant copy number variation from nucleotide 79,100 through 152,500 of chromosome 6, which determines resistance to DSM1.

## CONCLUSIONS

We present a robust tool to rapidly identify the genetic differences between any two related organisms. MinorityReport relates mapped sequence reads in SAM format from any alignment tool for both the mutant and parent genome, relative to a reference genome, and produces the set of variants that distinguishes the mutant from the parent. Output includes the spectrum of nonsynonymous changes, insertions or deletions, and copy number variations in a presumed mutant relative to its parent. Tunable evidence filters prioritize reported variants based on relative proportions of read counts supporting the genetic variant in the mutant versus parent data sets. Since MinorityReport is organism agnostic, it will be useful for both haploid and diploid genomes, as well as microorganisms and viruses.

The software is freely available, open source, and only requires Python with libraries included in standard distributions (re, math, sys, and os). We also include in the github distribution the R code for producing the data graphics shown in the Figures (this has software library dependencies), and a “master” script that breaks the calculation into smaller parts by chromosome and launches them in parallel for faster calculation of nonsynonymous variants on machines with more processor cores than there are chromosomes in the genome.

## LIST OF ABBREVIATIONS

SAM: Sequence Alignment / Map
GFF3: gene file format files
CNV: copy-number variant
SNP: single nucleotide polymorphism
PfATP4: sodium-dependent ATPase transporter
SRA: NCBI Short Read Archive
DSM1: dihydroorotate dehydrogenase inhibitor

## DECLARATIONS

### Ethics Approval And Consent To Participate

Not applicable.

### Consent For Publication

Not applicable.

### Availability Of Data And Material

All data are available on the NCBI Short Read Archive, and software is available as a supplement and will be deposited in GitHub upon publication. Data:<https://www.ncbi.nlm.nih.gov/bioproject/PRJNA253899/>

### Competing Interests

The authors declare that they have no competing interests.

### Funding

JAH was funded by NIH/NIDCR training grant T32-DE007306. WW was funded by the Philippine California Advanced Research Institutes project IHITM 63. JLD was funded by the Howard Hughes Medical Institute.

### Authors’ Contributions

JAH and JLD conceived of and designed the software and example applications. JAH wrote the software, prepared the figures, and drafted the manuscript. JLD and WW edited the manuscript and figures, tested and suggested improvements to the software. All authors read and approved the final manuscript.

## Acknowledgements

Not applicable.

---

**Supplementary Table 1.** Nonsynonymous variant data for a *Plasmodium falciparum* mutant strain raised for resistance to an apicoplast inhibitor, versus the susceptible parent strain. The causal mutation, IspD L244I, reaches near purity in the mutant, is absent in the parent, and receives the highest priority score.

**Supplementary Table 2.** Nonsynonymous variant data for a *Plasmodium falciparum* mutant strain raised for resistance to a dihydroorotate dehydrogenase inhibitor, versus the susceptible parent strain. Multiple mutations arise for all resistant strains, following the copy number variation and amplification shown in Figure 5.

